# Successful stimulation of myocardial ganglionic plexuses by TAU 20 in the absence of cardiac damage

**DOI:** 10.1101/2024.09.16.613202

**Authors:** Shengzhe Li, Jamie A Kay, Danya Agha-Jaffar, Cindy S Y Gao, Justin Perkins, Simos Koutsoftidis, Emm Mic Drakakis, Chris Cantwell, Liliang Wang, Prapa Kanagaratnam, Rasheda A Chowdhury

## Abstract

Atrial fibrillation (AF) is a major healthcare burden worldwide. For AF that is resistant to pharmacological intervention, the standard invasive treatment is a pulmonary vein isolation (PVI) procedure. Ganglionated plexuses (GP) ablation can be used as an adjunctive therapy to PVIs, together reducing the likelihood of AF recurrence. High-frequency stimulation (HFS) is a technique used to identify ectopy-triggering GP sites. However, to locate GP sites, sequential HFS must be delivered over the whole atria. Therefore, ensuring the safety of HFS delivery is integral to avoid causing irreversible damage from excessive pacing. We tested Tau-20 version 2 neural simulator, a prototype of a novel electrophysiological pacing and recording system that has the potential to guide intracardiac AF treatments. Using an *ex vivo* porcine Langendorff model that closely resembles the anatomy and physiology of a human heart, we confirmed that HFS can successfully trigger AF, indicating that HFS-positive locations contain GP sites. Additionally, we found that the HFS delivered via Tau-20 version 2 did not cause any damage to the heart. These findings evidence that once fully optimised, the Tau-20 system could be suitable for use in clinical settings.

## Introduction

Atrial fibrillation (AF) is one of the most common cardiac arrhythmias, affecting around 59.7 million people worldwide^1^. Over the last two decades, the incidence of AF has increased by 31% as the risk of AF rises with age^2^. AF leads to inefficient blood flow, and as such, a dangerous complication of AF is the formation of blood clots and stroke occurrence^3^.

When symptomatic AF is resistant to pharmacological interventions, pulmonary vein isolation (PVI) procedures^4^ have remained the cornerstone invasive techniques for AF treatment over the past 26 years^5^. However, permanent rhythm control is rarely achieved with PVI even with improvements to both technique and equipment. Moreover, it has been shown that the 2-year recurrence rate of AF is 5.8% and increases to 25.5% at 5 years^6^.

Recently, studies have shown that the intrinsic cardiac autonomic nervous system (ANS) plays a crucial role in initiating spontaneous AF^7,8^. ANS innervation to the heart occurs through an extensive epicardial neural network of autonomic ganglia, known as the ganglionated plexi (GP). The GP release neurotransmitters which can shorten the action potential and precipitate arrhythmia formation^9^. Evidence suggests that GP ablation in addition to PVI can improve the treatment success rate and reduce AF recurrence at 12 months post-intervention^10,11^.

High-frequency stimulation (HFS) is a technique used to locate GP sites. Sites positive for ectopy triggering (ET) or AF on HFS stimulation are considered sites of GP due to atrial arrhythmias being triggered at a rate faster than the local myocardial refractory periods^12^. An important factor to consider when sequentially mapping with HFS is how to deliver safe and effective stimuli.

Therefore, a prototype of a potential clinical pacing and mapping system (Tau 20 version 2) was tested in this study using an *ex vivo* porcine model to demonstrate that GP sites can be successfully stimulated with HFS without causing any damage to the tissue. Additionally, the proportions of tissue thickness and fat thickness were characterised to investigate the relationships between ET-GP sites and these factors.

## Results

Four porcine hearts were tested. HFS was delivered to 36 points of the atria. Four sites in two hearts demonstrated ET-positive locations. Twelve samples distant from the area of stimulation (>2mm) were also saved for histology as negative controls. Among all these four ET-positive sites, two were from the left atrium and two were from the right atrium. Detailed information of samples from each heart is shown in Table **??**. An example of the 3D model image of the hearts for electrode localisation is shown in Figure 1.

**Figure 1.**
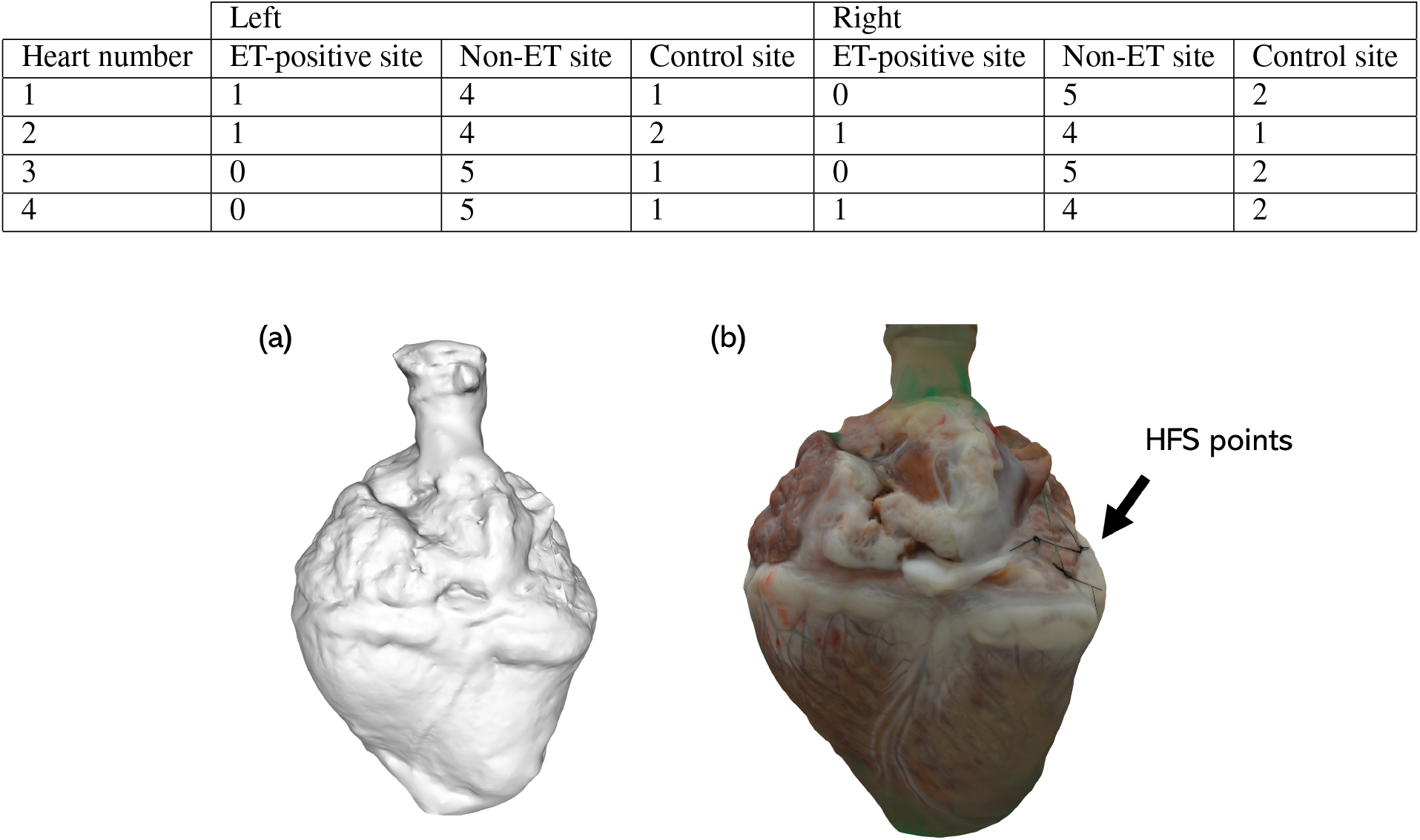
An example 3D image of a porcine heart. (a) A grayscale model; (b) A 3D model with real colour. The sites that had HFS delivered were labelled by suture. This is indicated by a black arrow on the image.

### Atrial fibrillation triggered by high-frequency stimulation

Sinus rhythm and HFS were recorded from all locations before and during the delivery of HFS. From the 32 negative sites, the EGM reverted to sinus rhythm immediately after the HFS and from the 4 ET-positive sites, AF was captured after the HFS. Examples are shown in Figure 2.

**Figure 2.**
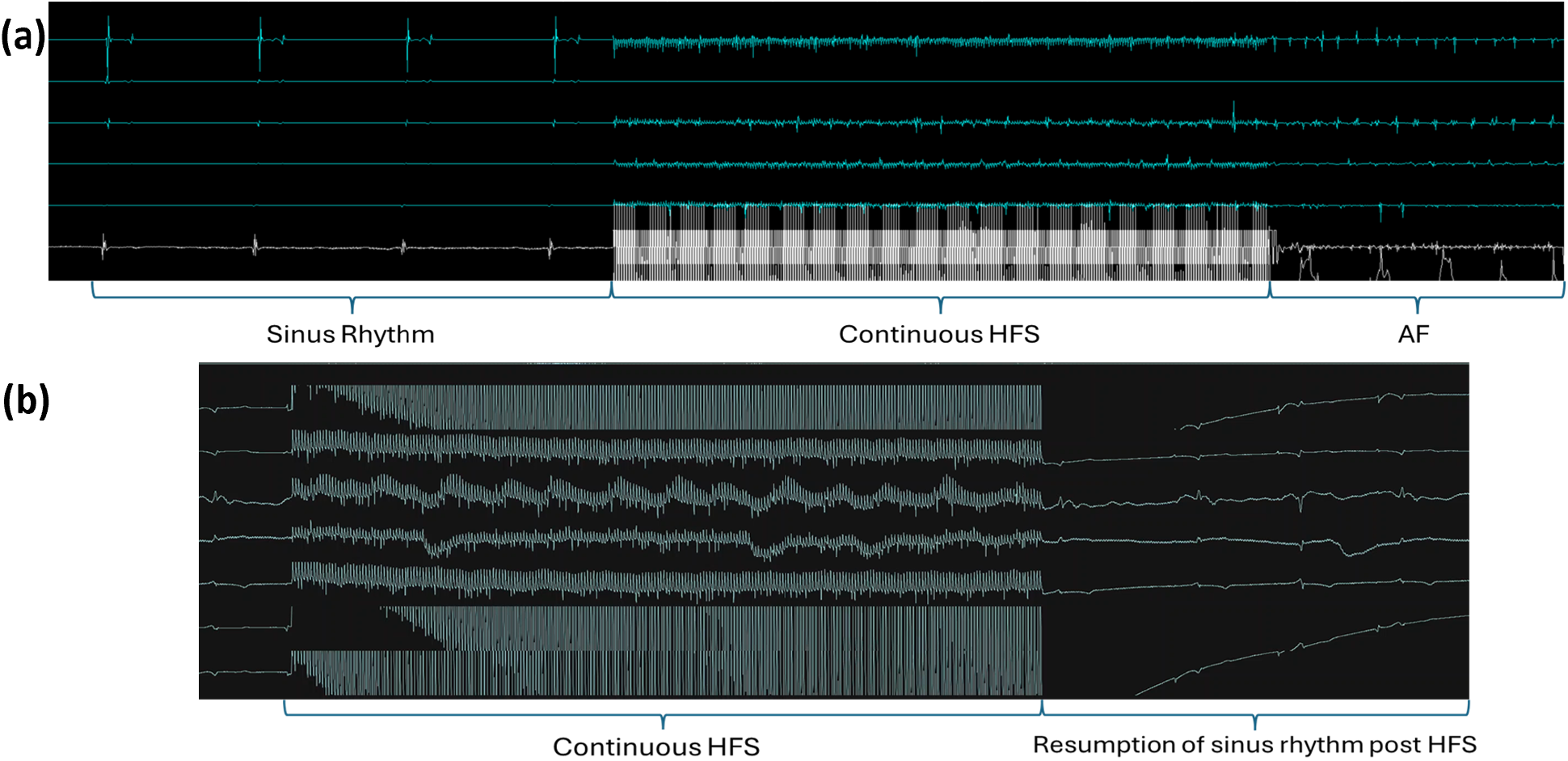
Examples of EGM recordings. (a) An example of an HFS-tested site triggering reproducible AF. Sinus rhythm was captured before the delivery of HFS and all the electrical activities of the heart were obscured by the HFS signal during its delivery. AF was captured after HFS and the same site was tested 2-3 times, consistently triggering AF in all instances.; (b) An example of a negative site. Same as the positive sites, sinus rhythm was captured before and after HFS and only the HFS signal was recorded during the stimulation.

### Damage level of the samples

To investigate whether damage was caused by the HFS delivered by the Tau 20 neural simulator, H&E staining was carried out on the samples from all the tissues (4 ET-positive sites, 32 ET-negative sites and 12 sites without HFS delivery) procured after the live protocols. At least two samples from each tissue were stained and the damage levels were quantified. There was no significant difference between the damage levels of ET-GP sites (mean value: 2.5, SD: 0.58), ET-negative sites (mean value: 2.4, SD: 0.50) and no HFS sites (mean value: 2.42, SD: 0.51) with a *p*-value of 0.93, 0.93 and 0.94 respectively. Examples of the H&E staining and group data are shown in Figure 3.

**Figure 3.**
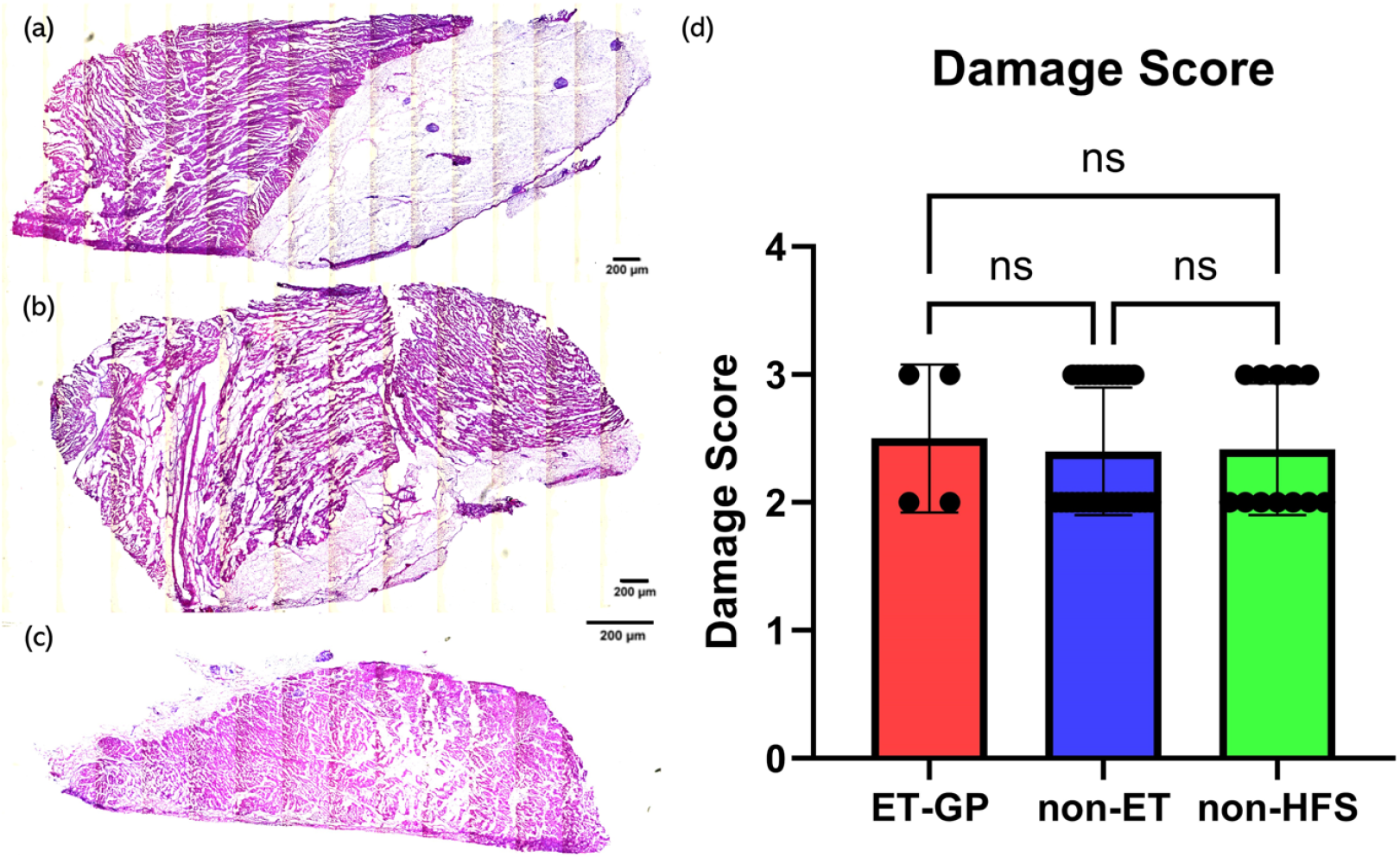
H&E staining examples and the damage score with relation to HFS. (a) An example of H&E staining of a sample from an ET-GP site.; (b) An example of H&E staining of a sample from a non-ET site.; (c) An example of H&E staining of a sample from a non-HFS site.; (d) Damage score with relation to HFS. Samples from 4 ET-GP sites, 32 non-ET sites and 12 non-HFS sites were analysed. At least 2 samples were scored from each tissue and the highest score was chosen to represent the damage level of the tissue. A one-way ANOVA statistical analysis was used to assess significance. ‘Remote’ refers to areas far away from compact fibrosis. ‘Near’ refers to areas close to compact fibrosis. The mean values are represented by each bar. The dots represent the individual value of each heart. Error bars represent the standard deviation. Statistical analysis results are shown on the top bracket where ns means not significant when *p*-value *≥* 0.05.

### Comparison of myocardial thickness and fat thickness between ET-positive sites, non-ET sites and control sites

To explore whether there is a correlation between myocardial thickness and fat thickness in ET-positive sites, non-ET sites, and control sites, the length of the total myocardial thickness, fat thickness, and their proportion from all the tissues (4 ET-positive sites, 32 ET-negative sites and 12 sites without HFS delivery) were compared. At least two samples from each tissue were measured. The mean value of lengths measured from the samples from the same tissue was calculated to represent the thickness of the whole tissue and the thickness of the fat. ET-GP sites (total thickness value: 3098.8um ±530.5um, fat thickness value: 1790.88um ±388.9um, mean percentage: 0.57 ± 0.03) demonstrated significantly thicker wall thickness, fat thickness and ratio compared to non-ET (total thickness value: 1500.23um 404.6um, fat thickness value: 267.65um ±296.8um, mean percentage: 0.15 ±0.15)/ non-HFS (total thickness value: 1563.54um ±366.2um, fat thickness value: 258.38um ±210.4um, mean percentage: 0.17 ±0.13) sites. No significant differences were observed between non-ET sites and non-HFS sites. Examples of the measurement and the result are shown in Figure 4.

**Figure 4.**
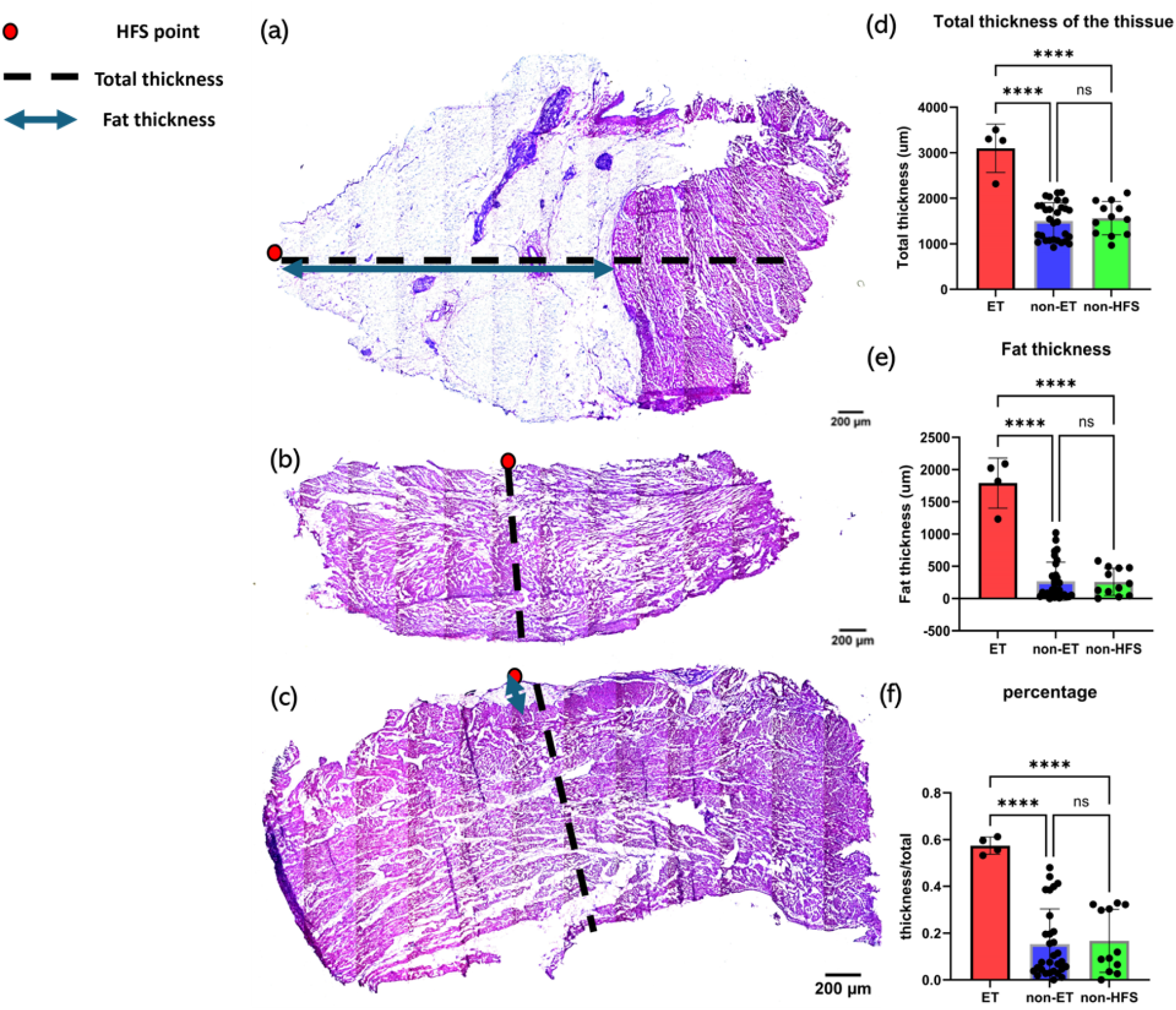
Examples of thickness measurements and thickness analysis. The red dot depicts the HFS positive site, the black dashed line represents the total thickness and the blue line represents the fat thickness. (a) An example of the thickness measurements of a sample from an ET-GP site.; (b) An example of the thickness measurements of a sample from a non-ET site.; (c) An example of the thickness measurements of a sample from a non-HFS site.; (d) tissue response to HFS as a function of total tissue thickness.; (e) tissue response to HFS as a function of the fat thickness of the tissue.; (f) tissue response to HFS as a function of the ratio of the fat thickness to the total thickness of the tissue. samples from 4 ET-GP sites, 32 non-ET sites and 12 non-HFS sites were analysed. At least 2 samples were measured from each tissue and the mean values were calculated to represent the thicknesses. A one-way ANOVA was used to assess statistical significance. A result was considered significant if the *p*-value ≤ 0.05. The number of asterisks represent different significance levels: * = *p*-value ≤ 0.05, ** = *p*-value ≤ 0.01, *** = *p*-value ≤ 0.001 and **** = *p*-value ≤ 0.0001.

## Discussion

We have demonstrated that HFS delivered by Tau 20 version 2 sufficiently triggered AF and localised GP sites without damaging the tissue. However, some damage was obtained from all the tissue samples irrespective of HFS stimulation. Therefore, these injuries were likely caused by the Langendorff perfusion. As the heart was placed on the Langendorff system for several hours, a slightly higher perfusion fluid pressure than *in vivo* conditions may have led to myocardial oedema and damage, injuries that have been illustrated in several studies^13–15^. However, the comparison between the negative control (non-HFS sites) demonstrates that the damage was not due to the HFS.

This study also illustrates that there is a clear link between tissue thickness and fat thickness with ET-GP locations. ET-GP sites are located in areas of greater wall thickness fat thickness and fat/tissue thickness ratio than non-ET sites. These findings could be explained by the fact that GP are typically found in fat. This suggests that GP are more likely to exist in areas with higher fat thickness, while areas with less fat do not have enough space for GP. Currently, the most common method of locating sites in clinical practice is the delivery of a large number of HFS across the atria, which is time-intensive and also risks triggering fibrillation if the HFS is not delivered properly^16–18^. Therefore, these novel correlations with wall thickness could provide another potential GP-locating method for treatments. If wall thickness is assessed prior to HFS protocols through non-invasive imaging techniques, HFS could be delivered only to the requisite areas. However, since only four porcine hearts were tested, further studies are required to establish the generalisability of this finding and also to verify if such a trend can be found in humans.

In conclusion, the Tau 20 shows promise as a novel platform for HFS stimulation to identify GP sites.

## Methods

All animal studies were ethically reviewed and carried out in accordance with ethical standards. The protocol was approved by the Royal Veterinary College (RVC) Animal Welfare and Ethical Review Board. All animals were housed and transported under conditions specified in the UK’s Animal Welfare Act 2006 and The Welfare of Farm Animals (England) Regulations 2007. Animal care, investigations, and euthanasia were performed according to the Animals (Scientific Procedures) Act 1986.

### Pig preparation and heart explanation

Four healthy 4-5-month-old white female pigs weighing 70-80 kg were rested for at least a week before the heart extraction to provide a stable heart for the study. Ketamine (20mg/kg) and Midazolam (0.5mg/kg) were injected intramuscularly (IM) for the purpose of premedication. Propofol (2-4 mg/kg) was used to induce anaesthesia via a 22 gauge intravenous catheter which was placed in the auricular vein (Carestastion 650, GE Healthcare, UK). Anaesthesia was maintained using inhaled sevoflurane (SevoFlo, Zoeitis, UK). Fentanyl (0.2 mcg/kg/min) was applied via infusion to achieve analgesia after an initial loading dose of 2mcg/kg. Multiple vital signs were monitored to ensure welfare and the quality of the heart. In detail, the blood pressure was measured using a catheter (Leadercath, Vygon, UK) which was placed in the femoral artery. Body temperature, electrocardiogram, capnography etc. were measured using a multiparameter monitor (Carestation 650, GE Healthcare, UK). The heart was flashed using cardioplegia during the procedure. An overdose of 0.7 mL/kg pentobarbital was applied to perform euthanasia at the end of the heart retrieval. The procedure was similar to our previous studies^19,20^.

### Explanted whole heart preparation and Langendorff

All hearts were perfused with cardioplegia solution (40mg/L) mixed with Heparin (12500I.U./L) before removal from the chest. All the hearts had a minimum of 3 cm of the ascending aorta remaining after extraction to allow cannulation for the Langendorff system. The extracted hearts were stored in ice-cold cardioplegia solution for transportation. The maximum cold ischaemic time was 90 minutes.

A custom-built Langendorff apparatus was constructed to keep the hearts alive for around 3-5 hours^19^. The apparatus consisted of a water heater to warm the solution, a two-layer 5L solution reservoir (custom supply from Radnoti Ltd.), heating water was pumped into the outer layer to heat up the solution, an oxygen supply to oxygenate the physiological solution to maximal saturation, a heating coil (custom supply from Radnoti Ltd.) to keep the physiological solution at the optimal temperature of 37±0.5 °C, a bubble trap (custom supply from Radnoti Ltd.) to remove the air bubbles in the solution to prevent potential damage to the hearts and a high flow peristaltic pump (Cole Parmer, UK) to circulate the solution around the system at a constant rate (90 rpm). The physiological solution was oxygenated warm Tyrode’s solution (10–3 moL/L: NaCl, 130; KCl, 4.05; MgCl2, 1.0; NaHCO3, 20; NaH2PO4, 1.0; glucose, 5.5; and CaCl2, 1.35; pH = 7.4). Tyrode’s solution was used to flash the cardioplegia solution, warm the heart and provide necessary glucose and ions for the heart to maintain the electrical activities and contractions while it beats. The solution chambers were connected to the aortic cannula and oxygenatedTyrode’s solution was perfused into the coronary arteries in a retrograde manner, thereby maintaining the metabolic, electrical, and contractile activity of the porcine hearts. The perfusate exited the coronary circulation into the right atrium and was subsequently expelled from the heart. On restarting, the heart was cardioverted using a Lifepak 20e defibrillator (Physio-Control, Inc., USA). ECG was recorded on Labchart (AD-Instruments, Oxford, UK) to monitor the condition of the heart. During stabilisation, the heart was paced from the basal region of the left ventricle at 10% above the threshold of activation (usually 2mV) using a clinical stimulator (Micropace EP Inc., USA). The High-Frequency Stimulation Protocol was carried out once the heart was stabilised (sinus rhythm was consistently obtained for 15 minutes). At the end of the live experiments, the heart was fixed with Tyrode’s solution-buffered formalin solution and stored in a 4°C cold room. The Langendorff setup is shown in Figure5.

**Figure 5.**
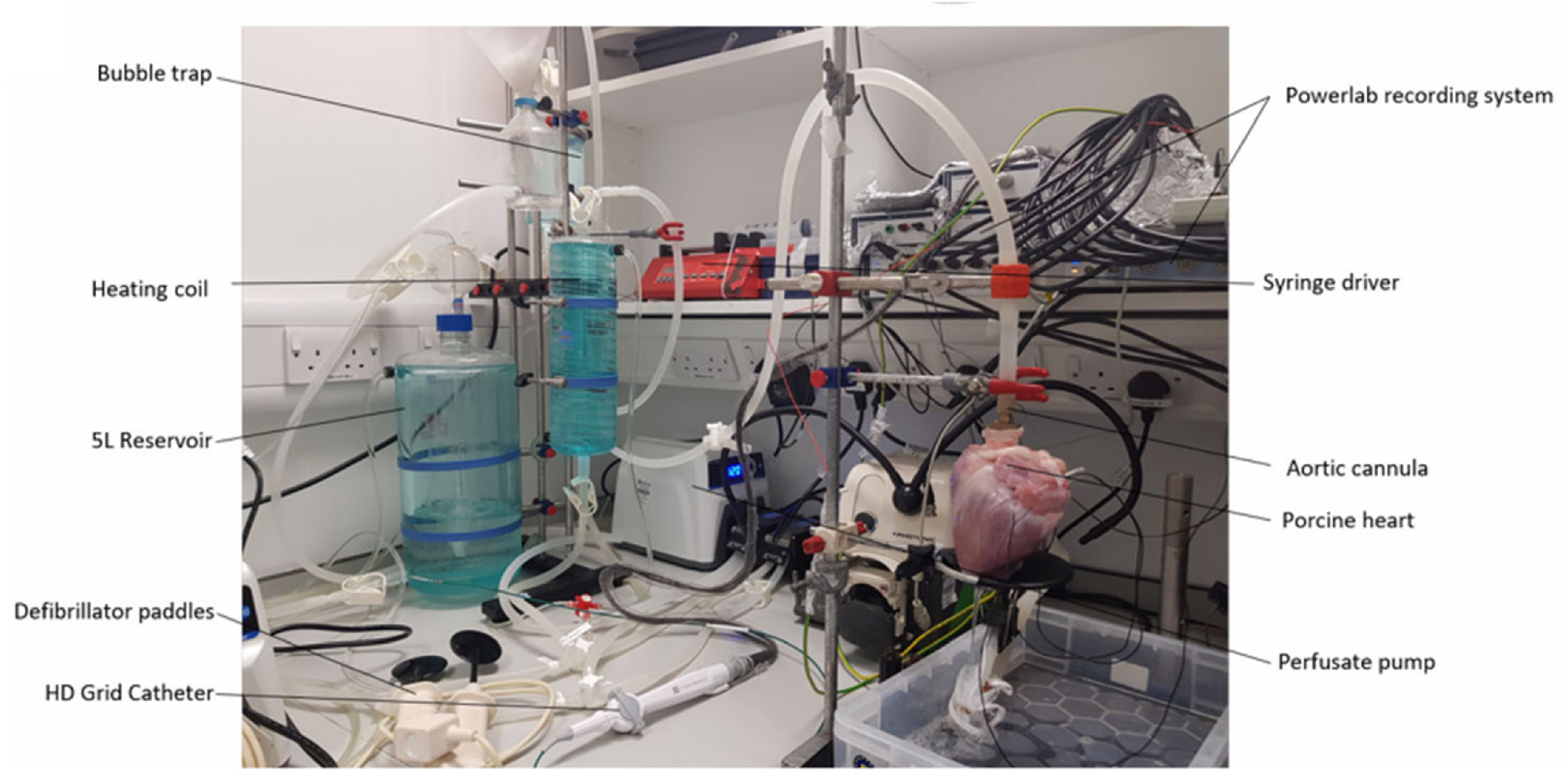
The Langendorff apparatus, pump, and aortic cannula feeding into a porcine heart.

### High-frequency stimulation

Tau-20 version 2, a prototype of a neural simulator was used in the study to generate HFS and record electrograms (EGMs). Tau 20 has similar functions to the Grass Stimulator which was commonly used in clinical studies and research. It can not be purchased or maintained in several countries due to the IEC/EN 60601-1 3rd Edition Regulatory Standards. The Tau-20 system is shown in Figure 6.

**Figure 6.**
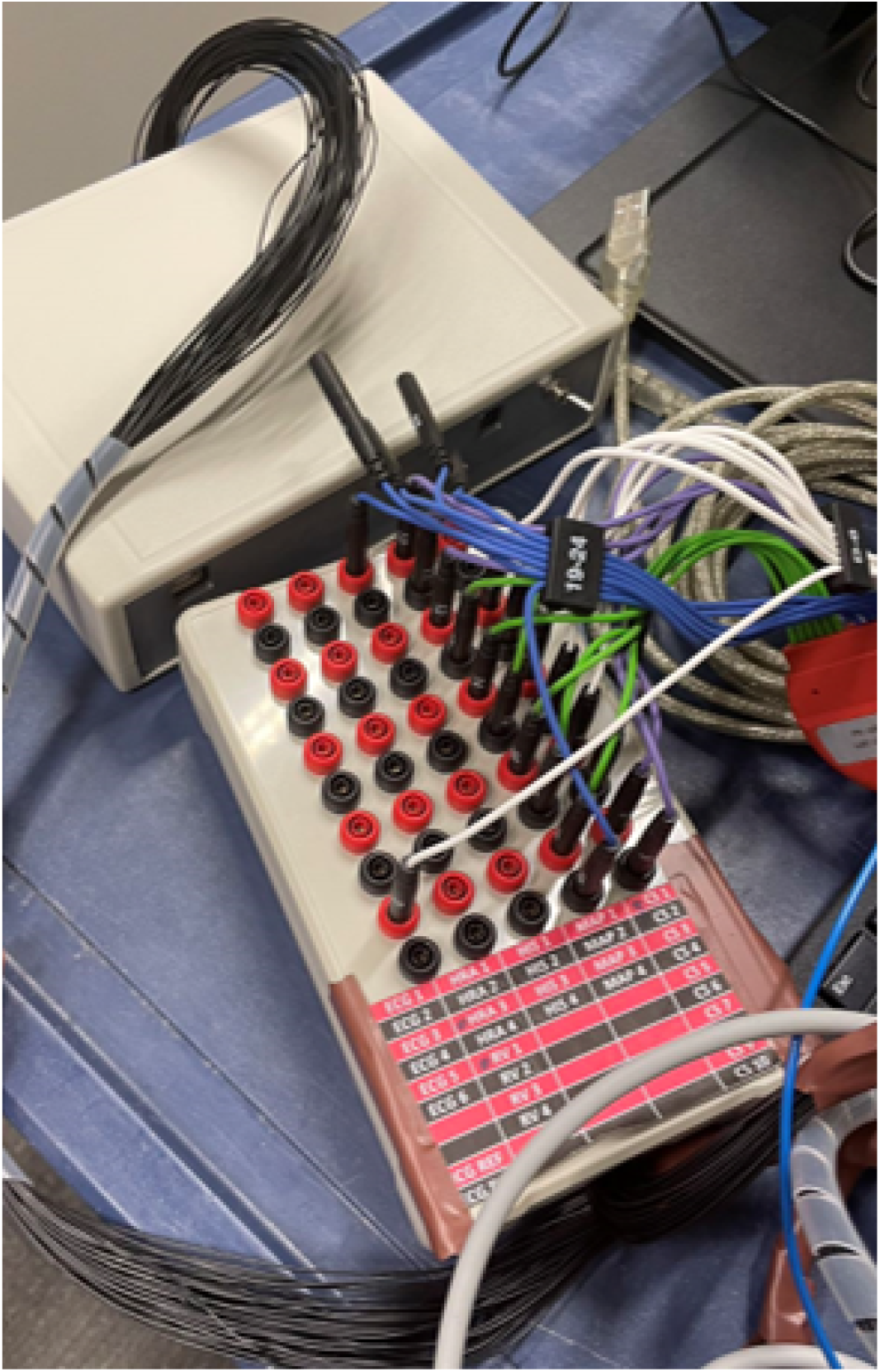
Tau-20 neural simulator and the breaking box for the catheter.

HFS was delivered to 6-10 locations on the epicardial surface of the right and left atria of each heart depending on size. HFS was delivered using a 4mm tip ablation catheter (Abbott, USA). In detail, the atria and the appendages were paced at high output (10v) at a fixed rate (600ms) to ensure that there was no ventricular capture. Once confirmed, HFS was delivered (pacing rate: 40 Hz, amplitude: 20mA, duration: 5 seconds). There are 15-20 mm distances between each stimulation point.EGMs were recorded next to the pacing points using an Inquiry™ AFocus II™ Double Loop Catheter (Abbott, USA) the whole time. A point was considered an ectopy-triggering(ET) site if a positive response was obtained from the real-time EGM recording after the HFS. Positive responses include atrial ectopy, atrial tachycardia, and AF. In most of the cases, it was AF. If no arrhythmia was captured, the stimulation point was considered a non-ET site. All the ET sites were tested at least twice to ensure reproducibility. All ET sites and non-ET sites were labelled with sutures to ensure they would be accurately dissected after the Langendorff for further imaging and staining.

### Histology of stimulated points

A tissue block (length, width:2mm, depth: from epicardium to endocardium) from each stimulated point was dissected from the heart once the heart was removed from the Langendorff and the whole heart image was taken before fixing the heart with formalin solution. The tissue block was wrapped with a layer of Parafilm and another layer of tin foil. The wrapped block was immediately put on dry ice for 15 minutes before being moved to −80°C freezer.

The frozen tissue block was then cut into sections (around 20 slices per block) with 10 µm thickness using an OTF 5000 Cryostat (Bright Instruments, UK), as cross-sections from epicardium to endocardium. The sections (2 on 1 slide) were then adsorbed onto the adhesion slides (Avantor, USA). The prepared slides were put back into the freezer for at least 24 hours.

Haematoxylin and eosin stain (H&E) staining was carried out using H-3502 Hematoxylin and Eosin Stain Kit (Vector Laboratories, USA) to test if the tissue was damaged by the HFS, H&E staining kit was used. The slides were fixed in ice-cold Acetone (Avantor, USA) for 20 minutes once they were taken from the freezer and dried off. The slides were stained with Hematoxylin for 30 seconds followed by two rounds of Milli-Q water washes (first 20 seconds, second 10 seconds). The slides were then stained with Blueing for 30 seconds and washed with Milli-Q water for 20 seconds. The final step of staining was 10 seconds of Eosin and the slides were dehydrated using Ethanol (Avantor, USA) after 20 seconds of Milli-Q water wash. The dehydration includes four steps: 70% Ethanol for 30 seconds, 90% Ethanol for 30 seconds and 2 rounds of 100% Ethanol for 30 seconds. After dehydration, the samples were cleared 3 times (5 minutes each) using Xylene (Honeywell, USA). The coverslips were applied with DPX mounting media (Sigma, USA).

H&E staining of the samples was used to visualise the damaged condition of the tissue. The concept utilised is a semi-quantitative scoring system used to measure the damage level of the tissue. The damage level score is shown in table1 Images were scored by eye, where a score of 1 was given for images with no damage and a score of 5 was given when the necrosis extends to the endocardial surface. If any sample exhibited multiple damage levels, the highest damage level was recorded.

**Table 1.** A semi-quantitative scoring system was used to assess the damage level. A score of 1 is given when there is no damage. A score of 5 is given when the necrosis extends to the endocardial surface.

| Damage Level | Damage Description |
| --- | --- |
| 1 | no damage |
| 2 | slight damage in the epicardial area |
| 3 | slight damage in the entire area |
| 4 | necrosis on the epicardial surface |
| 5 | necrosis on the endocardial surface |

### Imaging and analysis

The whole heart was imaged in 360 degrees using a Nikon camera from top, middle and bottom and these images were used to make a 3D model for the heart using ReCap Pro 2024 (Autodesk, USA).

The stained slides were imaged using a Zeiss Axio Observer inverted microscope controlled by ZEN Pro software (Zeiss, Germany) at Imperial College London. The objective used was 10x and the brightfield images were taken to examine the damage. The epicardial, myocardial, and connective tissue layers of each sample were determined by manual selection of the structures by histological examination, and their area was measured automatically using the software.

Image analysis was performed using ImageJ (FIJI) software. The thickness of the samples was recorded by measuring the length of the slides from the epicardium to the endocardium. The length of the fat layer was also measured to verify if there was any difference in fat and myocardial thicknesses between ET sites and non-ET sites. A semi-quantification system was applied to verify the damage level of the samples.

All images were blinded for analysis.

### Statistical analysis

GraphPad 10 (Prism, USA) was used for all statistical testing. One-way ANOVA analysis was performed to test for normality. The threshold for statistical significance was set to *p* < 0.05.

## Acknowledgements

The Langendorff apparatus used in this study was funded by the Rosetrees trust. The microscope used in this study was supplied by the Facility for Imaging by Light Microscopy of Imperial College London. Funding for this study was supported by the British Heart Foundation, welcome Trust and NIHR Imperial Biomedical Research Centre.

## Author contributions statement

Shengzhe Li, Jamie A Kay and Danya Agha-Jaffar conducted the experiments, Justin Perkins extracted the heart for the experiments, Shengzhe Li analysed the results. Liliang Wang, Chris Cantwell, Prapa Kanagaratnam and Rasheda Chowdhury oversaw the work, all authors reviewed the manuscript.

